# The California environmental DNA “CALeDNA” program

**DOI:** 10.1101/503383

**Authors:** Rachel S Meyer, Emily E Curd, Teia Schweizer, Zack Gold, Dannise Ruiz Ramos, Sabrina Shirazi, Gaurav Kandlikar, Wai-Yin Kwan, Meixi Lin, Amanda Freise, Jordan Moberg-Parker, Miroslava Munguia Ramos, Beth Shapiro, Jason P Sexton, Lenore Pipes, Ana Garcia Vedrenne, Maura Palacios Mejia, Emma L Aronson, Tiara Moore, Rasmus Nielsen, Harris Lewin, Paul Barber, Jeff Wall, Nathan Kraft, Robert K Wayne

**Affiliations:** University of California - Los Angeles; University of California - Merced; University of California - Santa Cruz; University of California - Davis; University of California - Berkeley; University of California - Riverside; University of California - San Francisco

## Abstract

Global change is leading to habitat shifts that threaten species persistence throughout California’s unique ecosystems. Baseline biodiversity data provide opportunities for ecosystems to be managed for community complexity and connectivity. In 2017, the University of California Conservation Genomics Consortium launched the California Environmental DNA (CALeDNA) program, a community science initiative monitoring California’s biodiversity through environmental DNA (eDNA)—DNA shed from organisms through fur, mucus, spores, pollen, etc. Community scientists collect soil and sediment samples, then researchers analyze the eDNA in the samples and share results with the public. The results are catalogues of thousands of organisms per sample, ranging from microbes to mammals. The CALeDNA website presents biodiversity inventories in a platform designed for the public and researchers alike, as well as user-friendly analysis tools and educational modules. Here, we present CALeDNA as a scalable community science framework that can harmonize with future biodiversity research and education initiatives.

## 1. Introduction

The Earth is facing unprecedented threats to its ecosystems due to climate change, habitat destruction, pollution and other anthropogenic factors. With the 6th mass extinction of life upon us (see Ceballos and Ehrlich, 2018), policymakers and the public need more information to address the grand challenges of how to protect, conserve and restore the health of vital ecosystems that provide food, medicines, raw materials, energy, and cultural attributes essential to human survival and well-being.

In California, one of three North American biodiversity hotspots (Myers et al., 2000; www.cepf.net), 40 million people must find a way to thrive while protecting biodiversity. The economy of California, now ranked fifth in the world, relies heavily on natural resources industries; the state ranks first in recreation tourism, second in seafood production, third in lumber production, and has 39 mined minerals that only occur in commercial quantities in our state (to learn more see www.conservation.ca.gov).

Inventories of California’s biodiversity are used to maintain these myriad ecosystem services residents rely on. However, detailed biodiversity data is hard to track across space and time. Fortunately, the past decade has witnessed an impressive rise in grassroots ‘community science’ (syn citizen science) campaigns to gather biodiversity data, such as through ‘<u>bioblitzes’</u> that monitor species presence or seasonal changes in organismal behavior, interactions, or development. While most community science initiatives are focused on gathering data, we argue that the state of California is a ‘living laboratory’ to testbed a feedback loop between the public and researchers, where all are engaged in data analysis and interpretation. With numerous world-class research institutions as well as curated living and *ex situ* natural history collections, and 18% of the U.S. colleges; hundreds of thousands of people in California already engage with environmental sciences and research (www.bls.gov). In addition, California has a strong naturalist certification program, created by the UC division of Agriculture and Natural Resources, where participation in community science is part of the curriculum.

The University of California (UC) Conservation Genomics Consortium (hereafter “the Consortium”) launched in 2016 with support from a UC President’s Research Catalyst Award. One aim, connecting research activities across campuses, was to develop a high throughput approach for community science-driven habitat monitoring and characterization using an environmental DNA method. In early 2017, the Consortium launched the statewide community science program called CALeDNA (*Cal ‘ee’ D-N-A*). CALeDNA is a platform for public and multi-institutional engagement in biodiversity data collection and analysis using DNA-based technologies executed in a series of steps (Figure 1). CALeDNA recruits and trains community scientists through its website, then coordinates soil and sediment collection using sampling kits and a phone app. Natural areas such as in the <u>UC Natural Reserve System</u> are sampled, analyzed for eDNA, and results are posted online in an interactive website and shared with natural areas managers and stakeholders.

**Figure 1.**
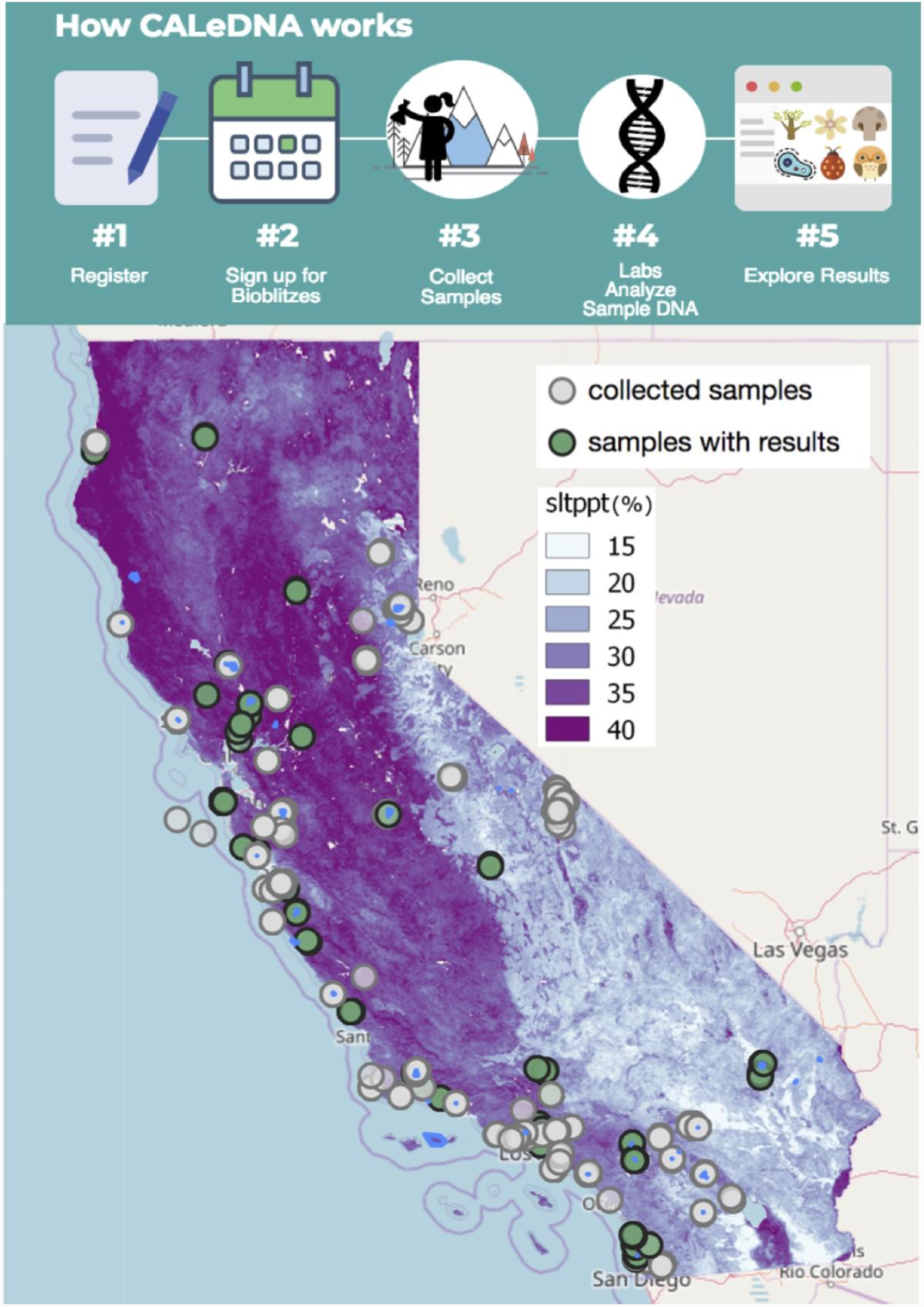
Top: Flowchart of how CALeDNA works. Bottom: Map of California showing the sites sampled by volunteers, and the proportion of samples for which eDNA results are publicly available.. Blue spots indicate the locations of UC Natural Reserves. Results from different organismal groups can be queried on the www.ucedna.com ‘explore data’ pages and plotted against different maps (example here shown is the proportion of silt in soils). The intention is for the user to do qualitative data exploration and generate hypotheses based on spatial patterns.

Diverse communities of researchers and the public have helped develop the research questions and the functionality of CALeDNA. Several California institutions with their own community science programs have partnered to organize bioblitzes and plan research projects (see section 4). Now, the program is focused on building an inclusive network with land managers, policy informers, naturalists, students, and university research scientists, as people are coming together to participate in the analysis of the open results and use the information to address grand challenges of how to steward ecosystems.

## 2. eDNA: the new biodiversity monitoring tool?

Environmental DNA is a promising solution to the challenge of monitoring marine, terrestrial and freshwater ecosystems (Bohmann et al. 2014; Thomsen and Willerslev 2015). eDNA survey methods rely on all organisms shedding DNA as they live and decay, and these DNA molecules can be isolated, sequenced, and identified (Taberlet et al. 2012). An eDNA-based inventory of a location is a kind of forensic reconstruction of the local organismal community (Thomsen and Willerslev 2015). DNA persists in surface soils and shallow sediments for variable lengths of time (mere days in the ocean, Lafferty et al., 2018; weeks or even several years in terrestrial environments, Barnes and Turner 2016). In all ecosystems, temperature, UV light, microbial metabolic activity, and eDNA shedding rates play complex roles in the production, movement, and degradation rates of eDNA (Barnes and Turner 2016; Deiner et al. 2017). Under certain conditions, like the bottom of a lake, eDNA may be protected from these physical and chemical threats, and may also be sheltered from consumption by active microorganisms (Palchevskiy and Finkel, 2006), leading to its persistence for up to thousands of years (e.g., Graham et al., 2016).

Next generation (high-throughput) sequencing technologies, such as Illumina MiSeq, HiSeq or NextSeq systems, substantially reduce the cost of DNA sequence data and allow thousands of different sequences to be retrieved simultaneously. This enabled the emergence of DNA <u>‘metabarcoding’</u>, in which specific DNA regions from any organism can be targeted, sequenced, and matched to reference <u>DNA barcodes</u> that communities around the globe have generated from <u>voucher specimens</u> for over three decades. Different barcoding regions are better for different constellations of organisms, but multiple regions can be targeted with metabarcoding, allowing a simultaneous inventory of biodiversity across organismal kingdoms, for costs currently as low as $35 a sample, and likely less in the future, as we optimize third generation sequencing technologies, such as PacBio (in progress). For CALeDNA, 4-6 regions are used to obtain metabarcodes from each sample, yielding lists of well over 1000 unique taxa per sample, representing all kingdoms of life. eDNA approaches are ideally suited for intensive and taxonomically broad biodiversity monitoring programs, where they may complement traditional field surveys, such as programs to test the impacts of global and local stressors on California ecosystems (Bohmann et al. 2014; Thomsen and Willerslev 2015).

The promise of eDNA monitoring has led to widespread development and application of this technique including large scale biodiversity monitoring networks (GEOBON and MBON), federal monitoring agencies (USGS and NOAA), local agencies (SCCWRP www.sccwrp.org), and research institutions (NHMLA). California’s research communities have pioneered DNA-based environmental assessments (e.g., Southern Sierra Nevada Critical Zone Observatory and the Aronson lab, see Aciego et al., 2017; Stanford Center for Ocean Solutions, see Andruszkiewicz et al., 2018). Diverse researchers and resource managers have been using eDNA approaches to detect and monitor endangered species, track the emergence and spread of invasive species, and inventory biodiversity in a wide variety of habitats from submarine canyons to alpine forests demonstrating the breadth of applications of this emerging technique. Work thus far has still largely focused on water sampling or focused on limited groups of taxa such as bacteria or fish (as in above two references).

## 3. CALeDNA program orientation

### 3.1. Study sites

Study areas can be chosen in two ways: (1) by researchers with projects, who propose collection in certain areas, habitat types, or transects, and who may organize group eDNA collection events, or (2) by community science volunteer choice. Volunteers can collect for CALeDNA from anywhere they please as long as they have proper permission such as collection permits or written permission from a landowner. While obtaining permission to collect eDNA may take time, it has not discouraged volunteers interested in adding an area of their interest to the CALeDNA map (Figure 1). CALeDNA reimburses all permitting fees incurred. This can also benefit groups, for example, one volunteer—a teacher—independently obtained a permit for Vasona Lake Park in Summer 2018, and brought the Youth Science Institute summer camp students to collect. Overall, the contribution of sites by the public and by researchers ensures a diverse sampling, increases awareness of accessible natural areas for all parties, and strives for sufficient sampling to meet research needs that will result in publications.

At the time of writing this, one third of our samples are from UC Natural Reserves. The UC boasts the largest university reserve system in the world, at 39 (soon 40) reserves totaling over 756,000 acres. Most of these reserves aren’t open to the public. UC researchers may visit, accompany volunteers, or even just send volunteers, to hike through and sample eDNA. The reserves are ideal to provide a biodiversity baseline for the state because they include coastal to montane biomes.

All reserves have hosted numerous traditional biodiversity surveys, and we use these to assess the extent of overlap between eDNA metabarcoding and traditional sampling, which can illuminate the bias as well as complementarity in eDNA and human surveys. The reserves offer additional abiotic data that may strengthen statistical analyses and models to describe eDNA patterns. These include weather station and tower data, such as that implemented by Institute for the Study of Ecological and Evolutionary Climate Impacts (https://iseeci.ucnrs.org), and <u>NASA pre-HyspIRI</u> flights, where for 7 years, data have been collected from pathways intentionally situated over UC reserves.

### 3.2. The sampling experience

Volunteers may join a bioblitz, or may sample a site on their own. In either case, they would receive a sampling kit of gloves, tubes, and an optional meter for collecting abiotic data (Figure 2a). Each sampling kit is used together with an electronic webform for smartphones and tablets or with a paper form. Forms are for the collector to provide important collection metadata (Figure 2b). These metadata fields are more than the minimum information currently required for meeting sample description standards (e.g. NCBI Bioproject), but additional data make samples more likely to be used for analysis. CALeDNA data standards are inspired by the Global Genome Biodiversity Network (ggbn.org).

**Figure 2.**
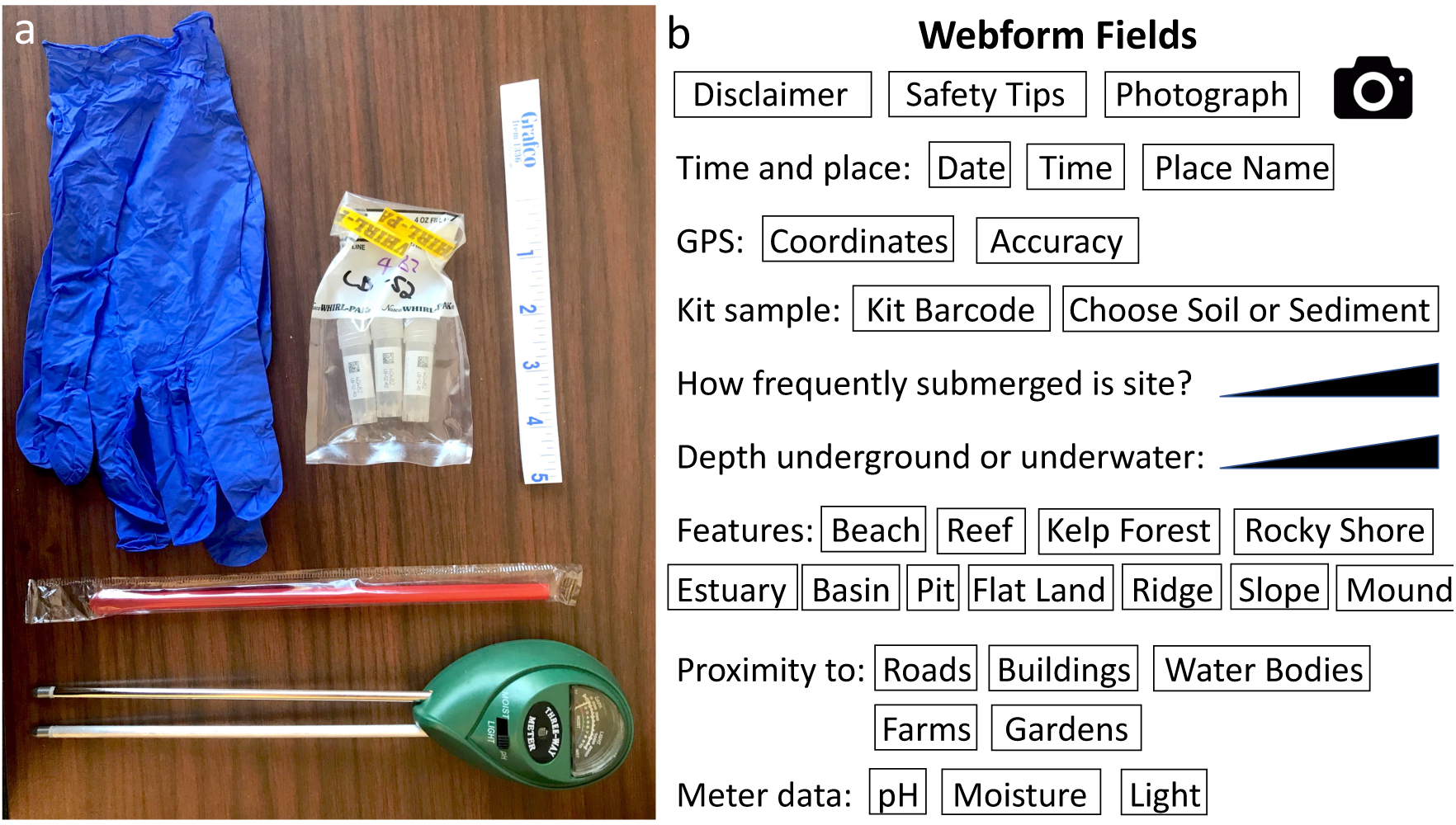
a. CALeDNA kit contents, including a pair of gloves, a set of three tubes for biological replicates packed inside a Whirl-Pak bag to protect tubes, a straw to sample sediment or to move large debris to expose topsoil, a ruler, and a meter. b. Webform fields the collector fills in when sampling a site.

Our webforms are made using the KoBoToolbox (kobotoolbox.org) platform to create and curate webform information. Results are backed up in real time. CALeDNA is dynamic, and different projects may require different metadata. Kobo Toolbox allows multiple forms to be created with the same minimum essential questions.

### 3.3. The ‘eDNA museum’

Upon receipt of the collected samples, each eDNA sample tube is treated as a valuable biological research collection. Samples get archived into a −80o C freezer that is part of the permanent “Dickey Collection” at UCLA, or archived in freezers at other UC campuses as satellite collections. We intend for the CALeDNA samples to be used to track environmental change over the next hundred years. When samples are processed and results are published online, the physical locations of the archived samples are reported and archived as part of the sample metadata.

Samples and kit materials are physically returned to UC campuses via pick up, drop off or Fedex. For the latter, we email shipping labels to volunteers so they do not need to pay out of pocket.

We encourage sample return within one week of collection. Many volunteers collect samples over long treks; in these cases, we request they refrigerate samples (4°C) until they can be shipped back all at once for archiving in our freezers. Tests have shown that freezing and thawing samples causes DNA profiles to vary, but maintaining a stable temperature helps to preserve the balance of DNA profiles (www.earthmicrobiome.org; Thompson et al., 2017). Considering the rapid advancement in technology, and our hopes that these eDNA samples will be used in research far in to the future, we chose to avoid adding stabilizing buffers to the samples that may pose unknown effects to the sample integrity.

### 3.4. Sample collection and laboratory processing

CALeDNA staff and interns continuously generate DNA data as sample collections increase. Under current funding, we are sequencing 10% of the samples received and make these results immediately open to the public.

Sample collection involves collecting three tubes from a site; these are treated as biological replicates. These replicates are thawed on ice, and a subsample of soil or sediment from each is pooled into a single tube that is mixed and used for DNA extraction. As a dynamic program, sampling methods may diversify in the future. For example, the Aronson Lab (UCR) is engineering rollers as eDNA surface collectors, along with wearable passive eDNA samplers.

DNA is processed through a series of steps to generate metabarcoding libraries. Because contamination from the sample collector or from the lab is a common problem in eDNA research, sometimes field ‘blanks’ are collected, and when extracting DNA, a ‘blank’ sample is also extracted as every batch of samples are processed. The details of the DNA preparation pipeline and CALeDNA protocols can be found on our website (www.ucedna.com) in the “researchers” space [DOIs to protocols will be assigned upon acceptance]. Each barcode region we target requires three separate PCR reactions as ‘technical replicates’ that help reduce reaction bias in the results, meaning for 5 barcoding regions, there may be 18 reactions per sample. Metabarcode libraries are sequenced on a MiSeq machine that generates paired reads 2 x 300 base pairs long, meaning when put together, each sequence can be up to 600 base pairs. This allows us to use lengthier barcode regions such as a portion of the *CO1* marker (Leray et al., 2013) to inventory animals. We aim to sequence a minimum of 25,000 paired reads for each barcoding region for each sample.

DNA data are deposited in the National Center for Biotechnology Information (NCBI) Sequencing Read Archive. These DNA data are processed through a series of software in the *Anacapa Toolkit* (Curd et al., submitted; https://www.biorxiv.org/content/early/2018/12/07/488627) that was specifically developed for CALeDNA’s multilocus metabarcoding approach. The toolkit combines state-of-the-art methods and is flexible to handle many kinds of eDNA data. CALeDNA researchers coordinating with eDNA researchers from academic, non-profit (Code for Science and Society), and government spheres to help onboard new user groups to *Anacapa*, which create opportunities for data integration.

Results are a list of taxa and the number of sequences that matched each one in each sample. The taxa may be identified to the level of species or limited to a higher rank such as genus or family, depending on the completeness of DNA barcode reference databases and the number of diagnostic DNA bases for that particular organism. CALeDNA scientists are working to solve this issue in the Nielsen Lab at UC Berkeley, but even in despite of this caveat, plenty of biodiversity patterns can be gleaned from higher taxonomic levels, like family, or from sheer genetic diversity.

### 3.5. Open data and results

To allow users to track our progress once samples are received, we put the field data collected by the community scientist online shortly after data are received. To make our results open and accessible, the eDNA results are deposited online shortly after processing through *Anacapa* and removing contaminants. Our impetus for open data is that scientists around the world are increasingly committing to the <u>FAIR data principles (FORCE11.org)</u> of findability, accessibility, interoperability, and re-usability. However, because endangered species may more easily be poached with help of eDNA leads, the CALeDNA website omits the specific sites where IUCN redlisted species have been found.

The *Anacapa Toolkit* is linked with an interactive results analysis platform called *ranacapa* (Kandlikar et al., 2018) that allows users to execute the same first-pass biodiversity data analyses of research projects as professional community ecologists typically do, but the automation in *ranacapa* relieves users of the need to code or use advanced statistical software. Plots and statistics are produced with explanations aimed at the undergraduate level. This enables community science users to reproduce results CALeDNA reports on the website or in scientific journals. Because data and tools are shared early in the analysis stage, community scientists may make some discoveries first, report them to CALeDNA, and through this feedback loop, earn co-authorship on research publications while bringing attention to the biodiversity in areas they care about.

## 4. CALeDNA research project vignettes

### 4.1. The Pillar Point project: assessing overlap between eDNA and human observation

Our first bioblitz in early 2017 was in collaboration with the California Academy of Sciences (CAS) and the Los Angeles Natural History Museum (NHMLA) to explore a potential complementary trifecta for biodiversity monitoring: human observation (CAS), DNA barcode sequences from local species (NHMLA), and eDNA (CALeDNA). Since 2012, CAS has been running monthly bioblitzes at the Pillar Point Harbor tidepools and adjacent areas within Half Moon Bay (https://www.inaturalist.org/projects/intertidal-biodiversity-survey-at-pillar-point), which is why this area was chosen. eDNA provides complementary results to human observation (Figure 3; manuscript in preparation; https://data.ucedna.com/research_projects/pillar-point).

**Figure 3.**
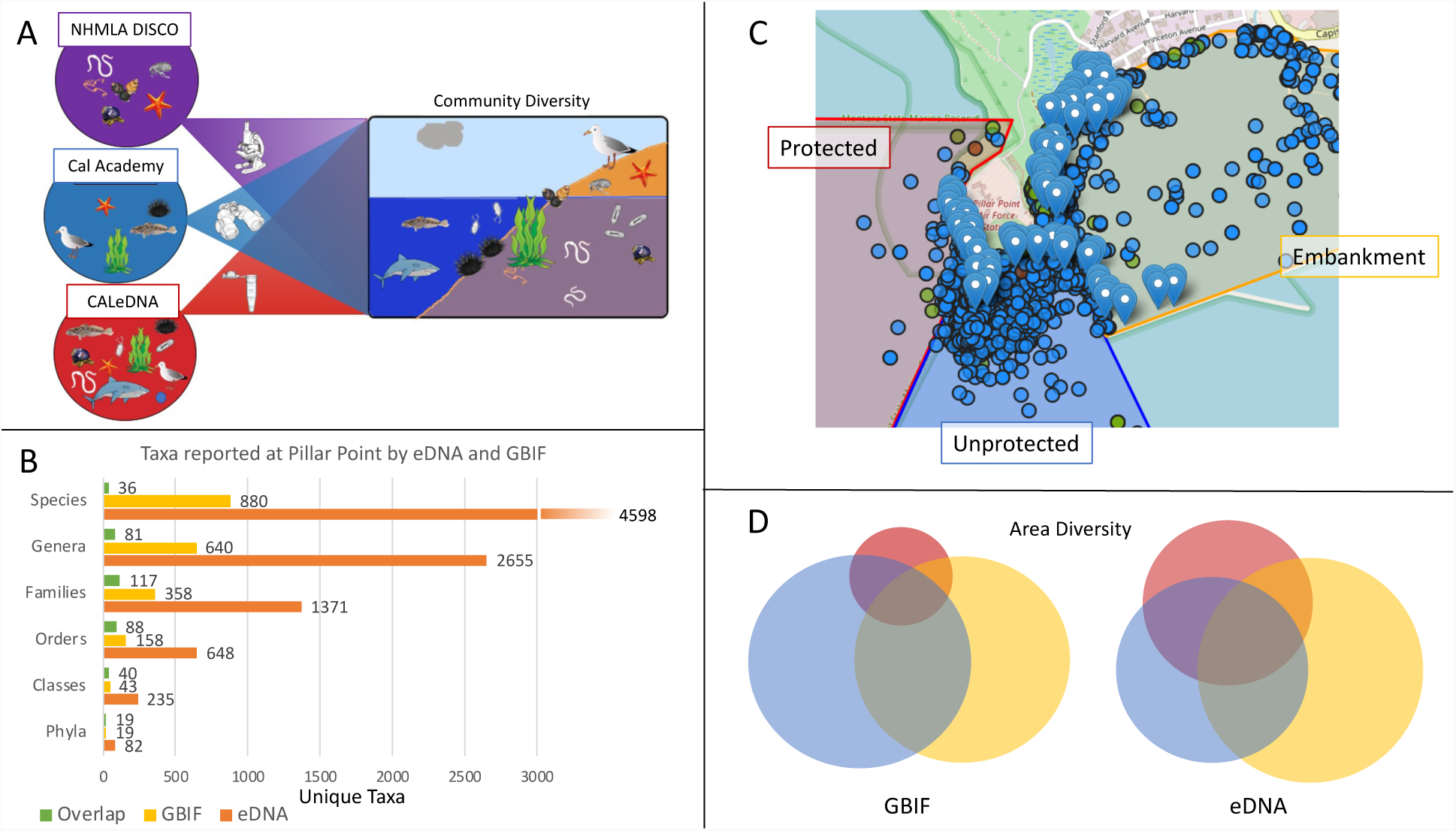
Pillar Point project overview. a. The project is an test of how observations, largely facilitated by the California Academy of Sciences iNaturalist program, integrate with local DNA barcoding efforts done by the Natural History Museum of Los Angeles Diversity Initiative for the Southern California Ocean (DISCO), and eDNA results from CALeDNA bioblitzes. These initiatives can cross-inform each other to broaden awareness of biodiversity that can be monitored through community science. b. Comparison of GBIF data, containing iNaturalist records and all other non-eBird observations, to eDNA. c. The Pillar Point project divides the region into three sections: an embankment (yellow), an unprotected exposed area containing accessible tidepools (blue), and the State Marine Protected Area (SMCA; red). The pins are eDNA sampling locations. The circles are GBIF observation records, colored by kingdom (blue is animal, green is plant, red is fungus). d. Area diversity showing the number of unique taxa observed from GBIF versus eDNA from the three sections of Pillar Point. Overlap is shared taxa. Colors for the sections are as in C.

### 4.2. Point Fermin: do eDNA results improve with local DNA barcoding?

NHMLA runs semi-annual bioblitzes in conjunction with Snapshot CalCoast (https://www.calacademy.org/calcoast) during low tide at Point Fermin Park in San Pedro, California (Figure 4a). They take photographs and make physical <u>voucher collections</u> as well, which later are DNA barcoded for the *CO1* region as part of the DISCO project https://research.nhm.org/disco/disco.html. eDNA collections concurrent with NHMLA bioblitzes help us assess how much results improve with very local DNA barcoding.

**Figure 4.**
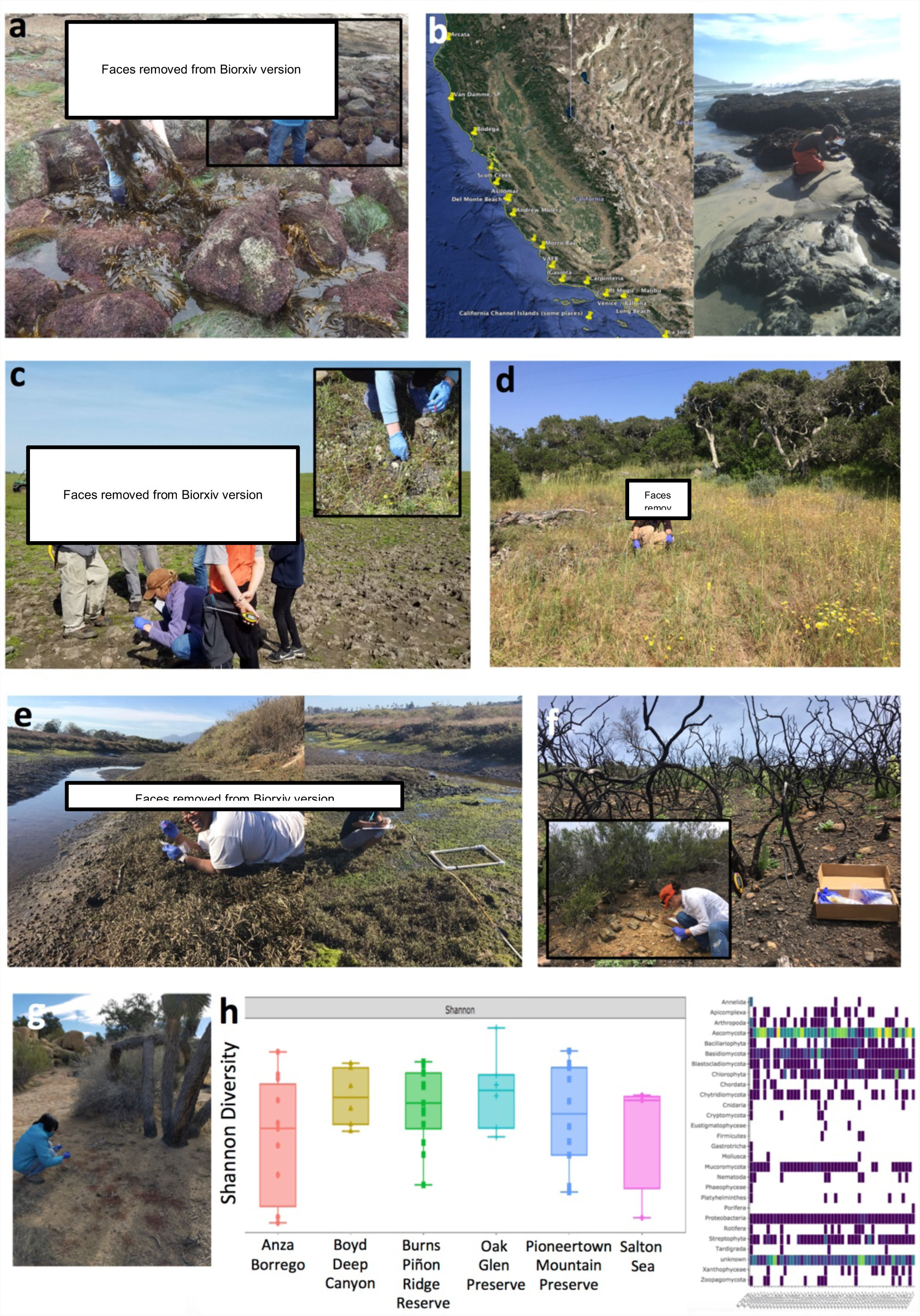
Project vignettes. a. NHMLA program coordinator [NAME OBSCURED] moves algae to uncover sediment for eDNA sampling by volunteers (inset). b. Left: the coastal bioblitz sampling scheme that occurs in the same weekend. Right: volunteer sampling the beach. c. Sampling the UC Merced Vernal Pool and Grassland Reserve. Biologists introduce their research to volunteers (here, [NAME OBSCURED], professor from CSULA, left, talks about fairy shrimp). d. Professors can be community scientists too: here [NAME OBSCURED], professor from CSUMB, hikes at UC Fort Ord Natural Reserve to collect for CALeDNA. e. [NAME OBSCURED] (left) samples along a lagoon. Volunteers (right) help count organisms using traditional ecology methods. f. Volunteer-submitted photos of paired burn samples from the Whittier Fire area. g. [NAME OBSCURED] sampling in the Mojave desert. She is now the CALeDNA web programmer. h. Left: Taxonomic richness is similar among the natural areas samples for the desert project. Oak Glen is a non-desert sample representative of DNA found in foothills that could wash into desert areas by runoff. Right: Presence of a taxon group (y-axis) across desert samples (x-axis). Variation prompts questions about ecological interactions among the stable members of the communities.

### 4.3. California macro-ecological patterns

From April 2017 to July 2017, a series of bioblitzes and independent community science activities in parks and reserves brought in thousands of soil or sediment samples to the CALeDNA collection. CALeDNA scientists selected 278 of these represented latitudinal transects along forest, shrub/scrub, or coastal areas down the state of California. Analysis of sequencing results reveals 25,283 unique taxonomic entries. We are performing different kinds of diversity analyses (e.g. Figure 5) and statistical modeling to ask what environmental factors influence biodiversity.

**Figure 5.**
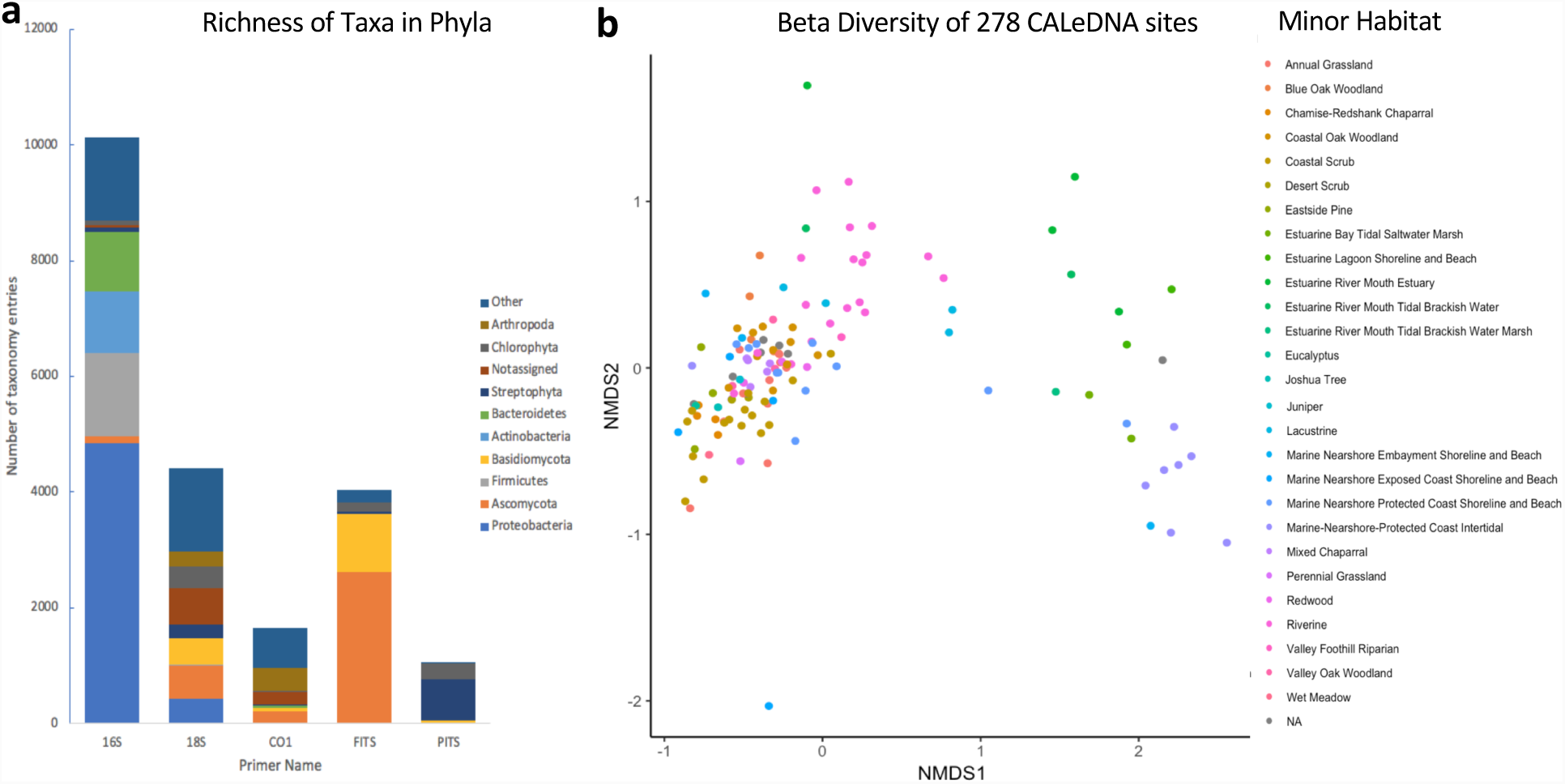
Taxonomic diversity plotted as a) unique taxon richness among phyla in results using different primers in metabarcoding to amplify different regions of DNA. 16S was chosen to amplify from bacteria and archaea. 18S was chosen to amplify the broad diversity of eukaryotes. CO1 was chosen to amplify DNA from animals. FITS (also called fungal ITS) was chosen to amplify all fungi. PITS (also called plant ITS2) was chosen to amplify DNA from angiosperms. The specific primers and methods used are on the CALeDNA website Methods for Researchers section: www.ucedna.com/methods-for-researchers. The ten most commonly found phyla were shown here. b. Non-metric multidimensional scaling plot (NMDS) showing beta diversity is similar for scrub and woodland habitats (left cluster), and these are very different from coastal samples (right). Each point represents one sample site, colored by the minor habitat it belong to. Habitat definitions from https://www.wildlife.ca.gov/Data/CWHR/Wildlife-Habitats.

### 4.4. Patterns of biodiversity along the California coast

Together with over two dozen colleagues from California State University campuses and coastal reserves, CALeDNA coordinated two distributed bioblitzes to sample along a 1200 km span of coast from Arcata to San Diego (Figure 4b). Over 80 phyla were identified and now, the team is asking how their presence predicts coastal health and uniqueness. These bioblitzes will be repeated to monitor coastal biodiversity change.

### 4.5. Persistence of eDNA in vernal pools

Vernal pools are temporary wetlands, filled by substantial rainy seasons, snowmelt, or groundwater. The pools host many California endemic species with special adaptations to pool depth, morphology and geochemistry. CALeDNA researchers from the UC Merced Dawson and Sexton labs are studying eDNA of five vernal pools on the UC Merced Vernal Pool and Grassland Reserve to build a more comprehensive taxon inventory (Figure 4c).

### 4.6. Invasive grasses in shrub/open forests

Invasive plants alter the community composition of fungi (Hawkes et al., 2006) plants (Gaertner et al., 2014) and microbiota (van der Putten et al. 2007) in the systems that they invade. The Fort Ord Natural Reserve has supported multi-day bioblitzes that have added nearly 200 samples to the CALeDNA collection with associated metadata of which sites have invasive grasses. UCSC graduate student Sabrina Shirazi is identifying associations between invasive species and the rest of the community detected with eDNA.

### 4.7. Biodiversity across lagoon systems

UC graduate students steer many CALeDNA research projects. Tiara Moore (UCLA; Fong Lab) brings community scientists to Carpinteria and Upper Newport Bay to sample sediment from different areas of lagoons (Figure 4e). She is evaluating the ability of eDNA to inventory community species and associate them with environmental stress response. DNA is being used in metabarcoding and also run on a GeoChip (Glomics, Inc) that quantifies the presence of 22,000+ genes involved in stress response and ecosystem functioning.

### 4.8. Burn sites

California has experienced and increase in fires and burn intensity that have devastated areas that are normally spared as refugia. CALeDNA community science volunteers and UC undergraduate classes began sampling paired burned and unburned sites (Figure 4f), and began resampling sites that were affected by fire. This will enable CALeDNA researchers to track biodiversity change after fire.

### 4.9. eDNA to describe the desert

UC Burns Piñon Ridge Reserve, Anza Borrego Reserve, and Pioneertown Mountain Preserve have hosted bioblitzes to help us understand the value of eDNA to detect a biodiversity in desert ecosystems (Figure 4g,h). Community scientists like [NAME OBSCURED] and Friends of the Desert Mountains contribute substantial collections for CALeDNA.

### 4.10. Exploring eDNA methods

The Shapiro lab at UCSC has tested how different approaches in preparing metabarcode libraries influence eDNA results that will help us tune methods to make CALeDNA research more efficient, low-cost, and have less technical bias. Past results have identified amplification enzymes that amplify DNA with less bias (Nichols et al., 2018). They continue to test technical effects on eDNA results.

## 5. eDNA in undergraduate education

### 5.1. Authentic research in the microbiology classroom

In Winter 2017, the newly launched CALeDNA initiative began a partnership with the UCLA Microbiology, Immunology, & Molecular Genetics (MIMG) department’s Course-based Undergraduate Research Experience (CURE) curriculum. CUREs have been demonstrated to provide a more inclusive avenue for students that might not otherwise have the opportunity to participate in research (Auchincloss et al. 2014). The MIMG CURE is a two-quarter research immersion curriculum in which upper-division undergraduates work in teams to formulate and test their own hypotheses regarding soil microbial ecology using eDNA and traditional bacterial cultivation methods (Shapiro et al. 2015). Using the CALeDNA sample collection kits and eDNA analysis tools, undergraduates have compared the soil microbiomes of California native and invasive plant species, natural and managed ecosystems, and studied the effects of human impact and burning on microbiomes.

Undergraduates connect with graduate students doing related eDNA research who visit the classrooms, and we hope this encourages students to consider graduate careers. This partnership between CALeDNA and MIMG inspired the development of eDNA and microbiology analysis tools spearheaded by graduate students and instructors, such as *ranacapa* (Kandlikar et al. 2018) and PUMA (Mitchell et al., in review; https://www.biorxiv.org/content/early/2018/11/29/482380). Several MIMG students have joined the CALeDNA labs as research interns.

### 5.2. eSIE: Environmental DNA for Science Investigation and Education

The Howard Hughes Medical Institute (HHMI) funded a novel project, **eSIE: Environmental DNA for Science Investigation and Education,** led by professors Wayne (UCLA) and Shapiro (UCSC). This program aims to educate and encourage undergraduates to enter STEM fields through field-based and <u>flipped learning courses</u>, workshops, and research, where eDNA gives entrée into the diverse natural and social sciences it can inform. An introductory course for freshmen and transfer students debuted in Fall 2018: *California’s DNA: A Field Course* (Figure 6, left). A 4-credit course, *Biodiversity in the Age of Humans*, is planned for Spring 2019 and will make use of the active learning classrooms at UCLA and UCSC campuses.

**Figure 6.**
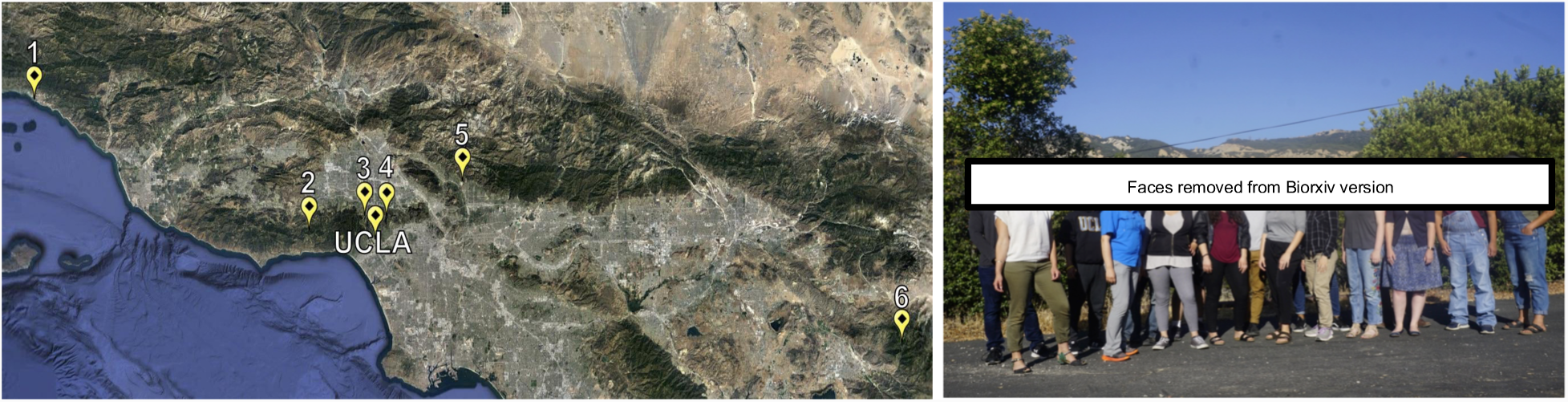
Left. The California’s DNA: A field course sampling locations for Fall 2018. Site number 1 is Carpinteria Salt Marsh Reserve, 2 is Stunt Ranch Reserve, 3 is the Skirball area that burned in 2017, 4 is Franklin Canyon Park, 5 is the Los Angeles River (Arroyo Seco), and 6 is the James San Jacinto Mountains reserve. The map was generated in Google Earth Pro. Right. Participants at the CALeDNA Summer Research Institute in Los Angeles. From left to right, [NAMES OBSCURED].

In Summer 2018, we launched two short-term *CALeDNA Summer Research Institute* sessions, in the Santa Monica Mountains (Figure 6), and in Santa Cruz, on the UCSC campus. The Institute was open to UCLA and UCSC undergraduates and extended to California State University, Los Angeles and Dominguez Hills. Activities were designed to prepare participants for beginning research projects in molecular labs. UCLA and UCSC offered eleven positions for 10-week paid summer research internships to work with 6 different professors after the Institute.

## 6. Building a stronger eDNA community

We hope to shatter the paradigms of the science that community scientists can do. We are continuously building resources for diverse user groups to use CALeDNA results and connect with university researchers through our web interface and our bioblitzes. A team of graduate student Information Architects as well an experiences web programmer with a passion for science were crucial to the production of the website. We encourage feedback and ideas for how to serve the community, and how to use eDNA science to inform policy.

In the next phase of the program we will tie CALeDNA into the Earth BioGenome Project (EBP; Lewin et al., 2018). The EBP is a moonshot to sequence the genomes of all eukaryotes on Earth. There are approximately 9000 eukaryotic taxonomic families on Earth (Lewin et al,. 2018), and at least 35,000 species in California. CALeDNA will provide information on where species are distributed and where new species may occur, so that those places may be sampled for the EBP collections. Our research teams are beginning to invent ways to use entire genomes to monitor demographic and evolutionary change with eDNA, not just occurrence.

The future will require a tremendous task force of CALeDNA community scientists, naturalists, observers, local scientific societies, biological collections and information curators, to help the EBP effort lead to solutions in California. Together, California can build a biodiversity-responsible and DNA-innovative economy to meet the challenges of climate change and a growing population.

## Acknowledgements

We thank the University of California Office of the President Catalyst Program (CA-16-376437), the Howard Hughes Medical Institute Grant (GT10483), NSF-DEB 1644641 (NK), NSF-DGE 1650604 (GK), NSF-GRFP 2015204395 (ZG). We thank Michael Dawson (UC Merced) for contributing to the CALeDNA program design, for coordinating the coastal bioblitzes, and for constructive comments on the manuscript. We thank the following people who have guided the CALeDNA program: Regina Wetzer, Dean Pentcheff, and Adam Wall of the Natural History Museum Los Angeles, Eric Crandall (CSUMB), Alison Young and Rebecca Johnson of the California Academy of Sciences, Greg Suba (CNPS) and Joseph Miller (UCSC). We thank former interns Amber DeVries, Laura Rabichow, Larysa Bulbenko, Aoife Galvin, Audrey Mahinan, and Eric Beraut. We thank the UCLA UX Team for guiding the web development. We thank all UC Natural Reserve Managers who made CALeDNA collection possible and inspired research questions, and especially acknowledge the contribution of the late Don Canestro (UCSB).

Note: <u>Underlined</u> words in the main text are intended for a sidebar glossary throughout the paper.

Sidebar Glossary (*will appear in the order they are introduced in the text*):

## eDNA (environmental DNA)

DNA from environmental samples such as soil, air, surfaces, or water rather than directly from an organism. The DNA in the sample may be shed from a living or dead organism such as from skin cells, or from an entire organism that was collected as part of the sample, such as from a microbe. eDNA degrades over time as it is exposed to the elements, and so where and how long it can be detected depends on characteristics of the environment.

## Bioblitz

hands-on, educational and fun community science activities such as bird or wildflower surveys. They usually occur in a day and often contribute to biological research, monitoring projects, or research resources (e.g., iNaturalist).

## UC Natural Reserve System

A network of 39 (soon 40) natural reserves across California that total 756,000 acres of land, and 50 miles of coastal shoreland (ucnrs.org). The reserves function to save representatives of all of California’s ecosystems for research, education, and public service.

## DNA barcodes

Short DNA sequences of a region that varies in sequence among species and therefore can be used to match DNA to a species or strain. DNA barcodes are usually sequenced from voucher specimens.

## Metabarcoding

Sequencing a specific DNA barcode region of a genome from multiple organisms within a single sample. The many resulting sequences are matched to known DNA barcodes allowing variants to be assigned to identify species present.

## Polymerase Chain Reaction (PCR)

A technique used in molecular biology to make many copies of a region of DNA to allow for sequencing. It is performed by adding a mixture of enzymes, free nucleotides, buffers and primers to DNA, and then putting the mixture through a series of specific heating and cooling incubations. Primers are short sequences designed to flank the segment targeted for copying and sequencing.

## Voucher specimen

A whole organism or part thereof, such as a plant cutting for an herbarium specimen, that is preserved for scientific use and used as a reference to confirm identity.

## NASA pre-HyspIRI flights

Since 2012, NASA has flown planes over parts of California, with priority over UC natural reserves, to collect various kinds of remote sensing data that describe the abiotic and biotic features of the local environment at high resolution. These data inform the HyspIRI satellite design under plan to launch in 2020.

## FAIR Data Principles

Principles of minimum standards for digital science information distribution to benefit data providers and data consumers, both machine and human, that were set in 2014 <u>(FORCE11.org)</u>. Data should be <u>F</u>indable, <u>A</u>ccessible, <u>I</u>nteroperable, and <u>R</u>e-usable.

## Alpha diversity

the mean species diversity or taxonomic richness in a location.

## Beta diversity

a measure of diversity between areas, which helps describe diversity turnover at a regional scale. Beta diversity accounts for the number of taxa common to both areas and the number of unique taxa in each area. It describes the change in community composition from location to location.

## iNaturalist

A community platform for photographing, geotagging, and identifying organisms. iNaturalist is a phone app maintained by the California Academy of Sciences. To date, nearly 187,000 species have been observed in 15,000,000 observations by 1.1M people.

## Global Biodiversity Information Facility

A web-accessible database of all species observations and collections. It houses information for >1B species occurrence records. DNA data have only just begun to be included as an ‘observation’ of a species (UNITE; GBIF 2018).

## Flipped Learning Courses

Courses where content is learned via media at home and classroom time is used to carry out exercises that apply content.

